# Hyperbaric Oxygen-Induced acute lung injury: A Mouse Model Study on Pathogenesis and Recovery Dynamics

**DOI:** 10.1101/2024.02.23.581279

**Authors:** Shu Wang, Zhi Li, Guangxu Xu, Xiaochen Bao

## Abstract

Hyperbaric oxygen (HBO) therapy is extensively used to treat a number of ailments. Although oxygen is crucial for survival, too much oxygen can be harmful. Excessive oxygen inhalation in a short period of time can lead to injury, and the lung is one of the main target organs. Lung injury induced by hyperbaric oxygen is notably more severe than that caused by normobaric oxygen, yet systematic research on such injury and its regression is scarce. In this study, mice were exposed to 2 atmospheres absolute (ATA), ⩾ 95% oxygen for 2, 4, 6, 8 hours. Changes in lung histopathology, inflammation and expression of chemokines, vascular endothelial permeability, 8-OHdG and apoptotic cells were detected before and after the exposure. These parameters were also measured immediately, 12 hours, and 24 hours following 6 hours of exposure to 2 ATA of ⩾95% oxygen. The study indicates that lung damage from HBO is histologically characterized by bronchiolar and alveolar dilation, atelectasis, inflammatory cell infiltration, and hemorrhage. At 2 ATA with over 95% oxygen for 4 hours, there is a significant increase in pulmonary vascular permeability, as evidenced by elevated Evans blue scores (*p* = 0.02). After 6 hours of HBO exposure, there is a significant rise in pulmonary tissue pathology scores, 8-OHdG levels, and inflammatory and chemotactic factors (such as IL-6, CCL2, CCL3, CXCL5, and CXCL10), along with intercellular adhesion molecule 1 (ICAM1), vascular cell adhesion molecule 1(VCAM1). Moreover, it was observed that these markers continued to progress even after leaving the hyperoxic environment, peaking at 12 hours and starting to recover after 24 hours, suggesting that the peak of inflammatory lung damage occurs within 12 hours post-exposure, with recovery beginning at 24 hours. However, the content of Evans Blue, reflecting vascular endothelial damage, and ICAM1, VCAM1 remain significantly elevated 24 hours after leaving the hyperoxic environment, indicating that the pulmonary vasculature endothelium is the most sensitive to damage and the slowest to recover in HBO-induced lung injury. These findings provide evidence for the prevention and treatment of acute lung injury complications caused by HBO.

**NEW & NOTEWORTHY:** This study systematically observed the development and outcome changes of ALI induced by HBO. In lung injuries caused by high partial pressure of oxygen, the pulmonary vascular endothelial cells are the first to be damaged and the slowest to recover. A 6-hour exposure to 2 ATA, ⩾95% oxygen of hyperbaric oxygen primarily causes oxidative DNA damage and inflammatory responses without significant apoptosis. The lung injury progresses even after leaving the high-oxygen setting, with inflammation peaking at 12 hours post-exposure and beginning to subside after 24 hours.

## INTRODUCTION

Hyperbaric oxygen (HBO) therapy is commonly used to treat various medical conditions, including decompression illness (1), diabetic gangrene (2), osteoradionecrosis (3), stroke (4, 5), carbon monoxide poisoning (6), and acute coronary syndrome (7). HBO therapy involves placing the patient in a chamber, increasing the pressure within the chamber, and providing 100% oxygen for breathing. This delivers a higher amount of oxygen to the tissues100% oxygen for respiration. In this way, it is possible to, which deliver delivers a greatly increased partial pressure of oxygen to the tissues. Unfortunately, oxygen in high doses is potentially toxic to normally perfused tissue. Too much oxygen may be harmful, it leads lead to oxidative stress and the production of reactive oxygen species (ROS), which can cause cellular damage and death (8). Therefore, HBO therapy may be harmful to certain patients due to increased free oxygen radical damage.

One of the most common complications of HBO treatment is acute lung injury (ALI) (9, 10), which significantly limits its clinical application. Prolonged exposure to oxygen pressures up to 1 ATA (normobaric oxygen) results in widespread pulmonary damage with inflammation, accumulation of pleural fluid, respiratory failure, and death (11). When oxygen pressures exceed 1 ATA (HBO), the development of lung injury is accelerated and worsened. Studies have shown that exposure to higher pressures of oxygen without seizures leads to a different pattern of lung damage compared to lower pressures (12). Our previous has demonstrated increased expression of cytokines, cellular apoptosis and endothelial dysfunction, were significantly increased in lung injury induced by HBO exposure compared with room air controls, in rats (13) and mice (14). Excessive ROS leads to endothelial damage and inflammatory activation, resulting in increased expression of inflammatory cytokines and transcription factors in lung tissue (15), such as tumor necrosis factor (Tnf), interleukin 1 beta (Il1b), interleukin 6 (Il6), and interleukin 10 (Il10) (16-18). Additionally, ALI continues to progress even 24 hours after HBO exposure, and necroptosis may be involved in the pathology of ALI induced by hyperoxia (19). Currently, there is limited research on the duration of HBO exposure that leads to ALI and the recovery progress of ALI after HBO exposure. It is important to determine both the occurrence and the recovery process of ALI induced by hyperbaric oxygen, as this will provide a foundation for preventing ALI caused by HBO.

In this study, mice were exposed to a hyperbaric environment with over 95% oxygen at 2ATA for durations of 2, 4, 6, and 8 hours. We examined changes in lung damage, inflammatory responses, endothelial permeability, oxidative stress, and cell apoptosis after varying exposure times. Additionally, the changes in the corresponding indicators in mouse lung tissues at 0, 12, and 24 hours after removal from the high-oxygen environment were examined to clarify the recovery process of ALI caused by hyperbaric oxygen.

## MATERIALS AND METHODS

### Animals and ethical statement

7-8-week-old, male C57BL/6 mice were used in all experiments. Mice were housed in isolated cages and acclimatized to our housing facility for one week before experiments. All studies with mice were approved by the Naval Medical Center Animal Care and Use Committee (ethics number: NMC-202009).

### HBO treatment procedure

Animal experiments were divided into two parts

Part 1: 30 mice were randomly divided into 5 groups. The HBO groups were placed in a cylindrical hyperbaric oxygen chamber with pre-placed lime at the bottom (Yantai Hongyuan Oxygen Industry Co., Ltd., Shandong, China). The lime at the bottom was placed to minimize water vapor and CO_2_ accumulation. Before pressurization, 100% medical oxygen was flushed through the chamber for 10 minutes to replace the air. Then, slow pressurization was applied to reach 0.2 megapascals for 10 minutes. Oxygen concentration was continuously monitored and maintained at ⩾ 95%. The CO_2_ concentration was kept below 300 ppm. The condition of the mice was monitored by a camera located in the chamber and was recorded by a computer outside the chamber. After exposure to oxygen for 2, 4, 6, and 8 hours at 0.2 MPa, mice were slowly decompressed to normal atmospheric pressure within 10 minutes. The control group (air control group) was placed in the hyperbaric chamber at atmospheric pressure with air. After exposure, mice were immediately anesthetized by intraperitoneal injection of 1% pentobarbital sodium (0.1 g/kg body weight), the chest was exposed, and the lungs were collected after clearing the blood by perfusion with cold PBS via the right ventricle. (Figure 1A)

**Fig. 1.**
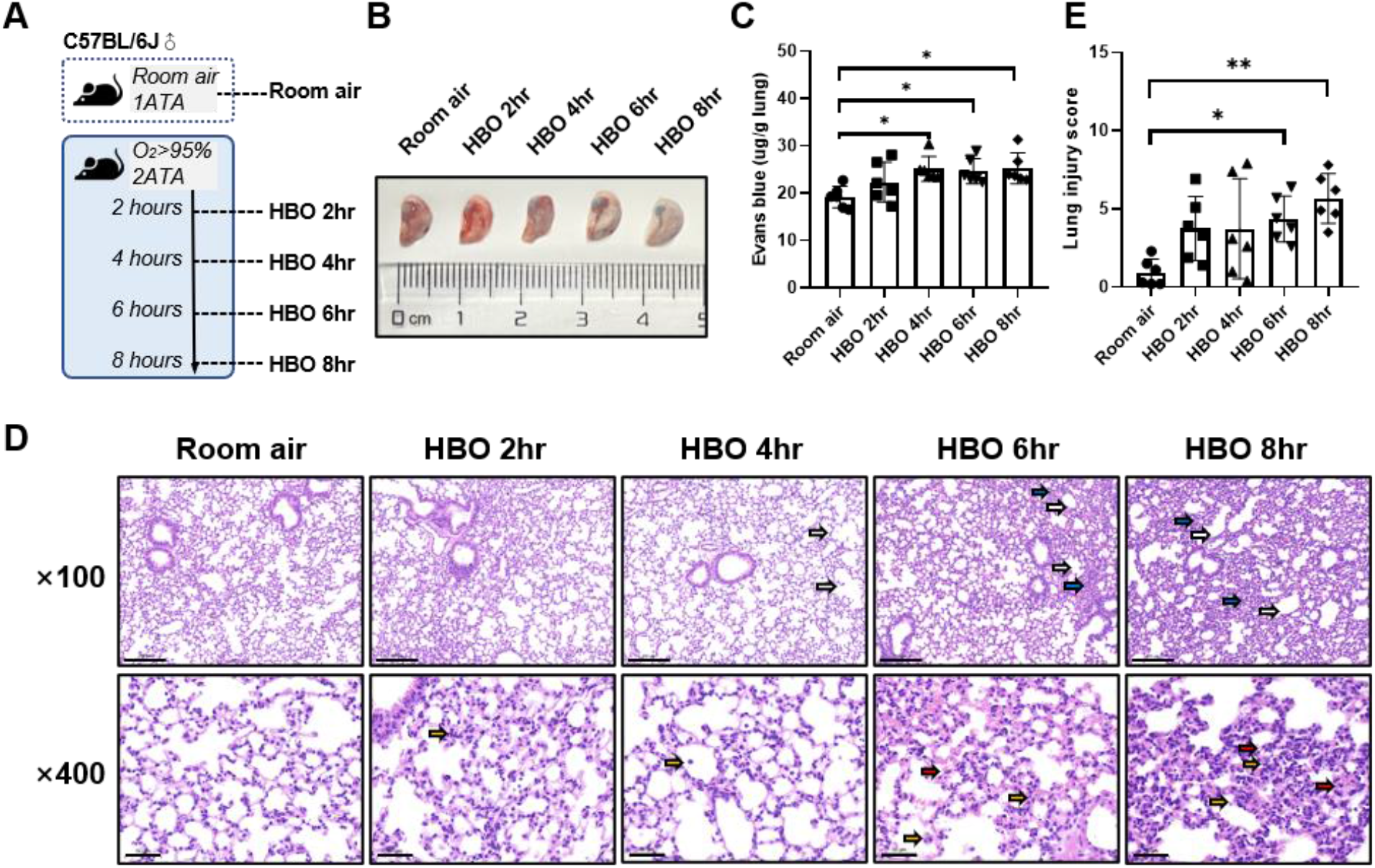
Lung histopathology and pulmonary vascular endothelial permeability in response to different durations of HBO exposure. (A) Experimental scheme. In the control group (Room air), mice were placed in the hyperbaric chamber at atmospheric pressure with air, while the experimental groups were exposed to 2ATA and ⩾95% hyperbaric oxygen for 2 hours (HBO 2hr), 4 hours (HBO 4hr), 6 hours (HBO 6hr), and 8 hours (HBO 8hr), respectively. (B) Representative image depicting extravasation of Evans-Blue dye in the lung. (C) Quantitative results showing the concentration of Evans-Blue dye in lung tissue. (D) Representative H&E-stained images of mouse lung sections. Yellow arrows: infiltration of alveolar and interstitial inflammatory cells; red arrows: alveolar and interstitial hemorrhage; blue arrows: atelectasis; white arrows: bronchial and alveolar dilation. Scale bars: 200 μm (100×) and 50 μm (400×). (E) Assessment of Smith lung injury scores for H&E-stained images. n=6, ^*^ p ⩽ 0.05, ^**^ p ⩽ 0.01, ^***^ p ⩽ 0.001.

Part 2: 24 mice were randomly divided into 4 groups. The control group (Control) was placed in the hyperbaric chamber under atmospheric pressure and exposed to ambient air. In HBO groups, mice were exposed to HBO for 6 hours and then depressurized to normal pressure. Lung tissue was collected at 0 hours (0 hours), 12 hours (12 hours), and 24 hours (24 hours) after exposure. (Figure 4A)

### Vascular permeability assay

Vascular permeability in the lungs was assessed using the Evans Blue (EB) assay. Mice were injected with 1% w/v Evans Blue solution at a dose of 40 mg/kg body weight intravenously, 45 minutes before sacrifice. To remove the intravascular dye, the mice were perfused with PBS through the right ventricle. The lungs were then flushed and homogenized using PBS (1 mL PBS/100 mg lungs). The EB was extracted from the homogenates by incubating and shaking overnight at 60 °C in formamide (twice the volume of the homogenates). After centrifugation at 5000r for 30 minutes, the concentration of EB in the lung supernatants was quantified using wavelength spectrophotometric analysis at 620 nm. A standard curve was generated by diluting the EB serially. The concentration of extravasated EB (μg of EB per g lung) in the lung was calculated based on the standard curve.

### Hematoxylin and eosin (H&E) staining and histological assessment of lung injury

For histological analysis, the lungs were fixed in 4% paraformaldehyde in PBS for 2 days, then paraffin-embedded and cut into 3 μm thick sections for H&E staining. From each section, 10 random areas were examined at a magnification of ×400. Within each field, lung injury was scored based on edema, inflammatory cell infiltration, hemorrhage, and atelectasis. Each stained feature was graded on a scale from 0 to 4: 0, absent and appears normal; 1, less than 25%; 2, 25 to less than 50%; 3, 50 to less than 75%; 4, 75% or higher. The four scores were added together and the total score used as the lung injury score for the sections. An investigator blinded to group assignment conducted all assessments.

### 8-hydroxyguanosine (8-OHdG) immunofluorescence

Immunofluorescence staining for 8-hydroxyguanosine (8-OHdG) was performed. Deparaffinized and rehydrated lung sections were incubated with FITC anti-DNA/RNA damage antibody (Ab183393, Abcam) at a 1:200 dilution overnight at 4 °C, followed by FITC goat anti-Mouse (green) at a 1:50 dilution. DAPI (G1012, Servicebio) was used for nuclear counterstaining. The numbers of total and positive cells were quantified using the Image J program (NIH, Bethesda, MD).

### TUNEL

TUNEL assay was performed on deparaffinized and rehydrated sections. The sections were stained with TMR (red) TUNEL apoptosis detection kit (G1502, Servicebio) following the manufacturer’s instructions. The nuclear counterstain was done with DAPI (G1012, Servicebio) in the dark. The number of total and TUNEL-positive cells per high power field was counted in 5-10 fields for each coded slide using the Image J program.

### Real-time quantitative polymerase chain reaction (qPCR)

Lung tissue total RNA was extracted in NucleoZOL (740404.200; Macherey-Nagel, German). The quality and integrity of extracted RNA were evaluated using Nanodrop Specthophotometer. The cDNA was synthesized using HiScript III All-in-one RT SuperMix. (R333-01, Vazyme, China). The housekeeping gene actin beta (Actb) was used as a reference. All primers used in this study were mRNA specific and designed for qPCR (Bio-Rad CFX Manager, USA) using ChamQ Universal SYBR qPCR Master Mix (Q711-02, Vazyme, China). Primers details are presented in Table 1. Melting-curve analysis was conducted for all PCR runs. Gene expression before and after the HBO treatment was compared using the 2−ΔΔCT (fold change) relative quantification method. The control group value and associated variability were expected to be close to 1.

**Table 1.**
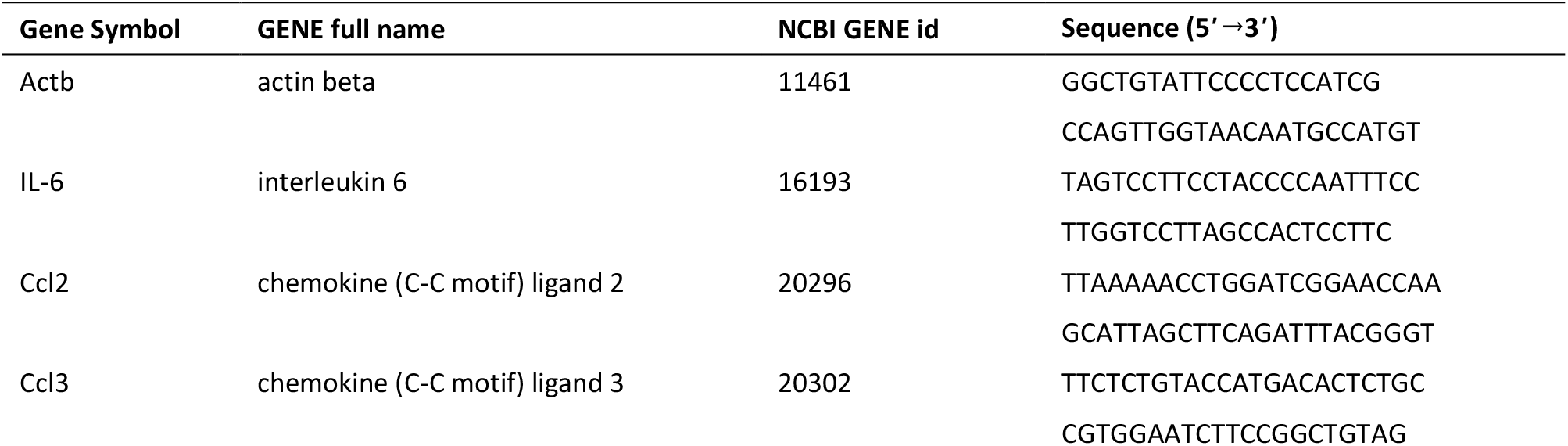
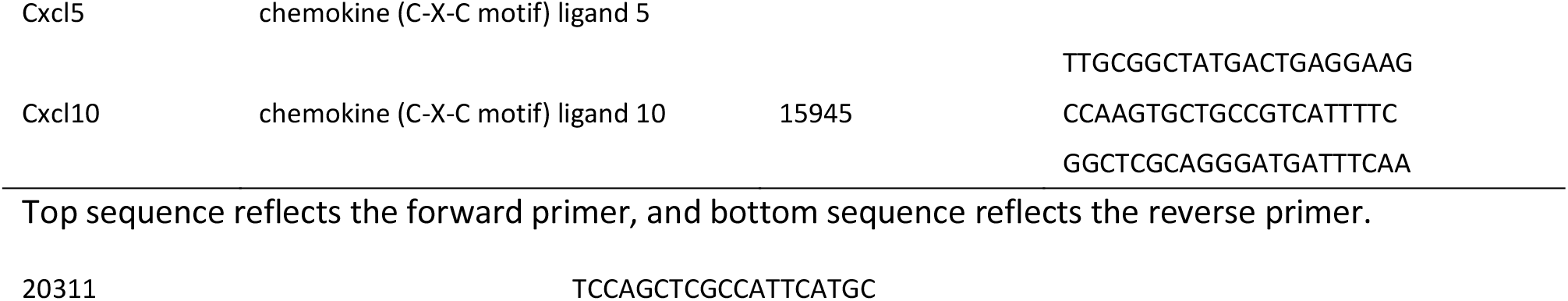
Nomenclature, gene information, and mRNA primer characteristics

### Western blot assay

Lung tissues were lysed using RIPA Lysis buffer (P0013B; Beyotime, China) containing protease and phosphatase inhibitor cocktail (dilution 1:100, 78442; Thermo Fisher Scientific, USA) for protein samples. The protein concentration was measured using the BCA method (23227; Thermo Fisher Scientific, USA). The protein samples were mixed with 5× sample buffer (MB01015; GenScript, US) and subjected to sodium dodecyl sulfate-polyacrylamide gel electrophoresis (SDS-PAGE). The electrophoresed protein samples were then transferred onto PVDF membranes. Following blocking with 5% fat-free milk for 1 hour at room temperature, the membranes were incubated overnight at 4°C with the following antibodies: ICAM1 (dilution 1:1000, ab222736; Abcam), VCAM1 (dilution 1:1000, ab134047; Abcam), and beta actin (dilution 1:5000, 20536-1-AP; Proteintech). Subsequently, the membranes were probed using Image Studio (Ver 5.2, LI-COR) after incubation with corresponding secondary antibodies (dilution 1:5000, SA00001-1, SA00001-2, Proteintech) for 1 hour at room temperature.

### Statistical analyses

Statistical analyses were performed with GraphPad Prism (GraphPad Software, La Jolla, CA). The results are presented as means ± standard deviation (SD). One-way analysis of variance (ANOVA) was used to compare the measurement data among groups. A *p* value less than 0.05 was considered statistically significant.

## RESULTS

### Lung histopathology and pulmonary vascular endothelial permeability in response to different durations of HBO exposure

Following HBO exposure, all mice survived without obvious abnormal symptoms. Focal bronchial and alveolar dilatation in lung tissues were observed in the 4-hour HBO exposure group using HE staining. In the 6-hour HBO exposure group, atrophy of the bronchi and alveoli adjacent to the expanded bronchi and alveoli was observed. Compared to the air control group, the 6-hour HBO-exposed group showed a significant increase in alveolar hemorrhage and inflammatory cell infiltration. Lung histopathology scores revealed significantly higher injury scores in the 6- and 8-hour HBO-exposed groups compared to the air control group (4.350 ± 1.464, 5.667 ± 1.586 vs. 0.933 ± 0.859; *p* = 0.0461, *p* = 0.0031) (Figure 1E).

EB staining was used to measure pulmonary vascular permeability. The results indicated a tendency of increased pulmonary vascular permeability with longer durations of HBO exposure, as depicted in Figures 1B and C. The 4-, 6-, and 8-hour HBO-exposed groups exhibited significantly increased pulmonary vascular permeability compared to the air control group (25.12 ± 2.628, 24.66 ± 2.661, 25.26 ± 3.268 vs. 19.17 ± 2.265 μg/g lungs; *p* = 0.0200, *p* = 0.0361, and *p* = 0.0166, respectively).

### Oxidative DNA damage and apoptosis in lung tissue in response to different durations of HBO exposure

Continuous in *vivo* HBO exposure leads to the accumulation of reactive oxygen species (ROS), resulting in the formation of 8-hydroxy-2’-deoxyguanosine (8-OHdG), a biomarker for secondary metabolites of oxidative DNA damage. Immunofluorescence analysis revealed a significant increase in the number of 8-OHdG-positive cells in the lung tissues of the 6- and 8-hour HBO groups compared to the air control group (3.606 ± 1.772, 4.640 ± 1.332 vs. 1.468 ± 0.802; *p* = 0.0406, *p* = 0.0012) (Figure 2A and B). Apoptosis in lung tissue, another important manifestation of HBO-induced ALI, was determined using the TUNEL immunofluorescence assay. The number of apoptotic cells in the HBO-exposed group was minimal and did not significantly differ from the control group (Figure 2C and D).

**Fig. 2.**
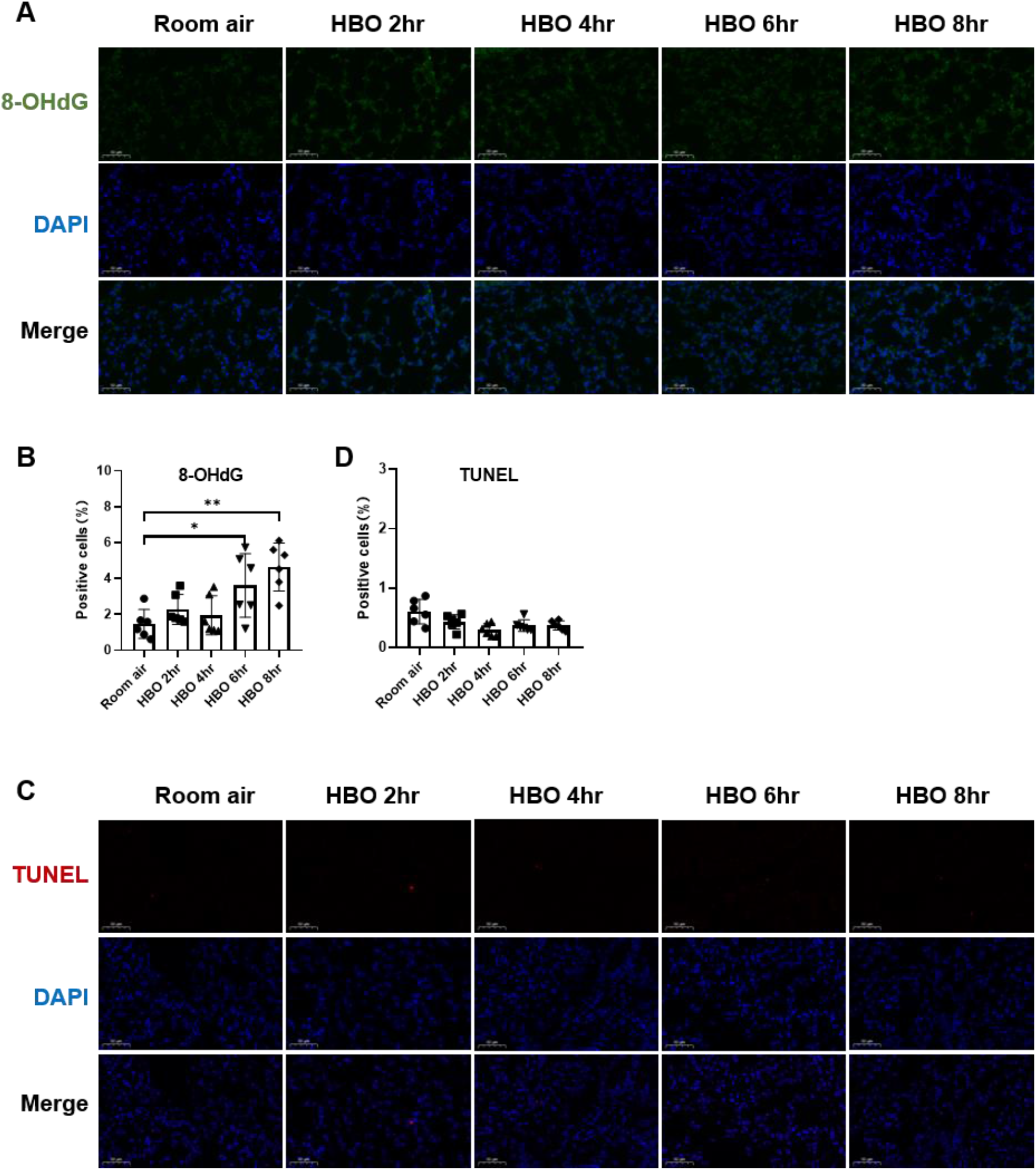
Oxidative DNA damage and apoptosis in lung tissue in response to different durations of HBO exposure. (A) Representative images of 8-OHdG staining to assess DNA oxidative damage in lung tissues. Scale bars: 50 μm. (B) Quantitative analysis of 8-OHdG in lung tissue sections as described in the Methods section. (C) Representative images of TdT-mediated dUTP Nick-End Labeling (TUNEL) staining showing apoptosis in lung tissues. Scale bars: 50 μm. (D) Quantitative analysis of TUNEL in lung tissue sections as described in the Methods section. n=6, ^*^ *p* ⩽ 0.05, ^**^ *p* ⩽ 0.01, ^***^ *p* ⩽ 0.001.

### Inflammatory factors and chemokines in lung tissues in response to different durations of HBO exposure

qPCR analysis was conducted to assess the expression of inflammation-related cytokines and chemokine genes in the lungs (Figure 3). The lung tissues of the 6-hour and 8-hour HBO groups exhibited significantly higher expression of IL-6 and Cxcl10 compared to the air control group (IL-6: 3.952 ± 1.036, 4.104 ± 1.129 vs. 1.000 ± 0.1392; *p* < 0.01; Cxcl10: 1.813 ± 0.4570, 1.959 ± 0.3410 vs. 1.000 ± 0.1407, *p* < 0.01) (Figure 3A and E). Ccl2 expression was significantly increased after 4 hours of HBO exposure (2.947 ± 1.152, 2.868 ± 0.7163, 4.229 ± 0.9360 vs. 1.000 ± 0.1019; *p* < 0.01) (Figure 3B). Ccl3 expression was significantly elevated in the 8-hour HBO group (1.201 ± 0.05853 vs. 1.000 ± 0.05024, *p*=0.0008) (Figure 3 C). However, HBO had no significant effect on the mRNA expression of Cxcl5 (Figure 3 D).

**Fig. 3.**
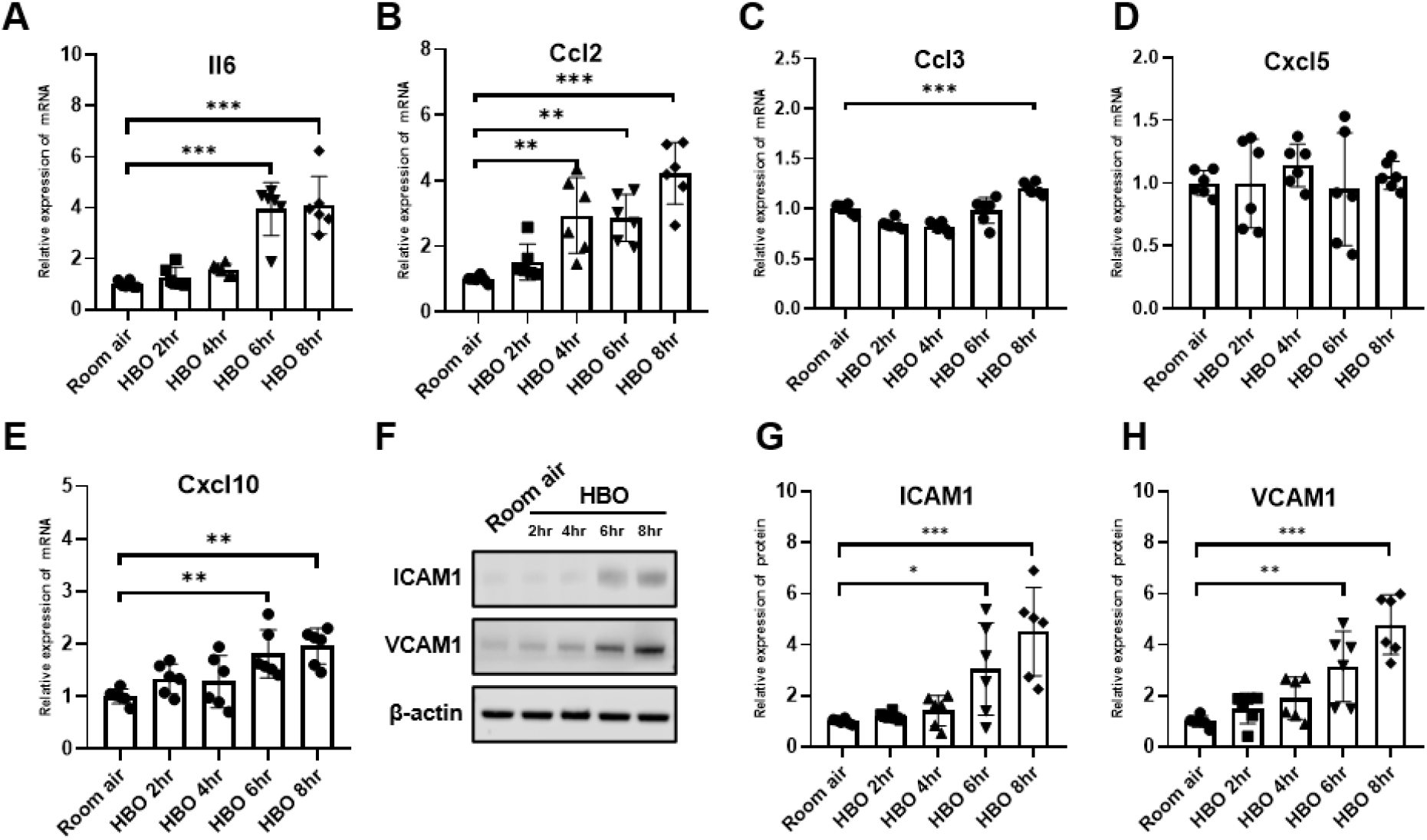
Inflammatory factors and chemokines in lung tissues in response to different durations of HBO exposure. (A-E) Expression of inflammatory cytokines and chemokines in lung tissues detected by qPCR as fold change. (F) Western blot analysis of ICAM1 and VCAM1 in mouse lung samples; beta actin used as the loading control. (G, H) Quantitative graph of western blot analysis. n=6, ^*^ *p* ⩽0.05, ^**^ *p* ⩽ 0.01, ^***^ *p* ⩽ 0.001.

Western blot was performed to assay for ICAM1 and VCAM1. The expression of both the ICAM1 and VCAM1 proteins increased with the extension of HBO exposure duration (Figure 3 F). In the 6- and 8-hour HBO groups, the expression of ICAM1 and VCAM1 were significantly higher than the air control group (ICAM1: 3.049 ± 1.815, 4.517 ± 1.729 vs. 1.000 ± 0.099; *p* < 0.01; VCAM1: 3.152 ± 1.379, 4.790 ± 1.167 vs. 1.000 ± 0.217; *p* < 0.01) (Figure 3 G and H). These data suggest that inflammatory response and endothelial dysfunction increased when exposed to HBO for more than 6 hours.

### Changes in histopathology and pulmonary vascular endothelial permeability in lung tissue during the recovery period after exposure to HBO

After 12 and 24 hours of 6-hour-HBO exposure, all mice still survived without obvious abnormal symptoms. Through HE staining of lung tissues, the dilatation and atrophy of bronchial and alveolar, alveolar hemorrhage, and inflammatory cell infiltration were still evident after 12 hours of 6-hour-HBO exposure, after 24 hours, these injuries improved. The lung injury score of the 0hr and 12hr after HBO groups were significantly higher than the Room air group (3.867 ± 1.148, 3.450 ± 0.841 vs. 0.917 ± 0.828; *p* = 0.0001, *p* = 0.0008, respectively), and the lung injury score of the 24hr after HBO group was significantly lower than the 0hr after HBO group (2.233 ± 0.894 vs. 3.867 ± 1.148; *p* = 0.0316) (figure4 D and E).

**Fig. 4.**
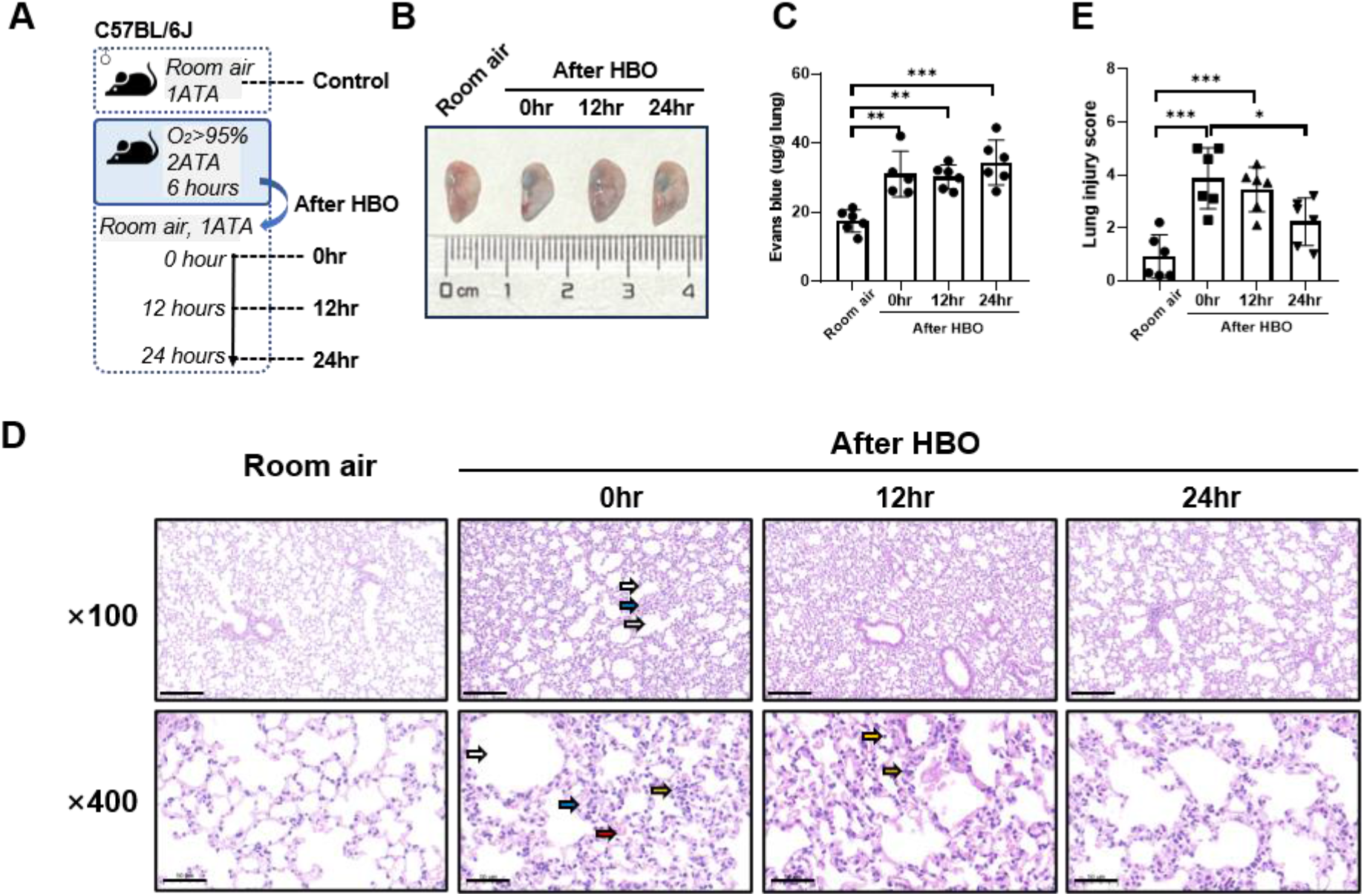
Histopathology and pulmonary vascular endothelial permeability in lung tissue during the recovery period after exposure to HBO. (A) Experimental scheme. In the control group (Room air), mice were placed in the hyperbaric chamber at atmospheric pressure with air, while the experimental groups were exposed to 2 ATA and ⩾ 95% oxygen for 6 hours. Lung tissue samples were collected from experimental groups at 0 hours (0 hr), 12 hours (12 hr), and 24 hours (24 hr) after the exposure. (B) Representative image depicting extravasation of Evans-Blue dye in the lung. (C) Quantitative results showing the concentration of Evans-Blue dye in lung tissue. (D) Representative H&E-stained images of mouse lung sections. Yellow arrows: infiltration of alveolar and interstitial inflammatory cells; red arrows: alveolar and interstitial hemorrhage; blue arrows: atelectasis; white arrows: bronchial and alveolar dilation. Scale bars: 200 μm (100×) and 50 μm (400×). (E) Assessment of Smith lung injury scores for H&E-stained images. n=6, ^*^ *p* ⩽0.05, ^**^ *p* ⩽0.01, ^***^ *p* ⩽0.001.

EB in lung levels showed less pronounced trend after HBO, as the figure4 B and C showed, compared with the Room air group, the EB in lungs of the 0hr, 12hr, and 24hr after HBO groups were significantly higher (31.06 ± 6.604, 30.21 ± 3.461, 34.41 ± 6.473 vs. 17.44 ± 3.220 μg/g lung; *p* = 0.0016, *p* = 0.0019, *p* < 0.001, respectively), but no significant difference between 0hr, 12hr, and 24hr.

### Changes in oxidative DNA damage and apoptosis in lung tissue during the recovery period after exposure to HBO

The expression levels of 8-OHdG and the number of apoptotic cells in lung tissue were measured during the recovery period following exposure to HBO. The results showed that the number of 8-OHdG-positive cells in lung tissue was significantly higher at 0 hours and 12 hours after removal from the high oxygen environment compared to the air control group (3.679 ± 1.593, 3.613 ± 2.291 vs 0.567 ± 0.301, *p* = 0.0122, *p* = 0.0144), but there was no significant difference between 0 hours, 12 hours, and 24 hours (Figure 5 A and B). TUNEL staining showed that there were still very few apoptotic cells in lung tissue at each time point during the recovery period, and there was no significant difference compared to the air control group (Figure 5 C and D).

**Fig. 5.**
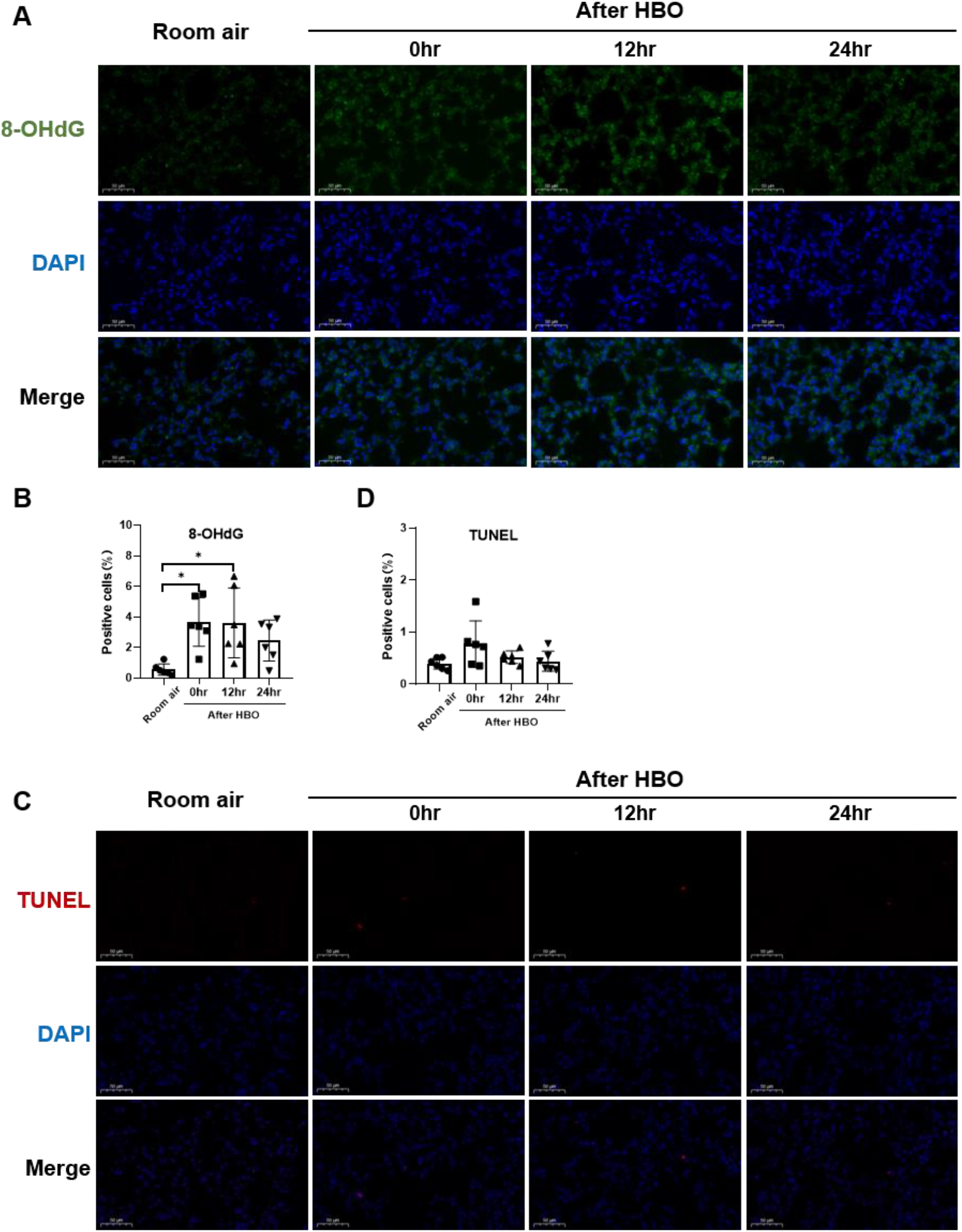
Oxidative DNA damage and apoptosis in lung tissue during the recovery period after exposure to HBO. (A) Representative images of 8-OHdG staining to assess DNA oxidative damage in lung tissues. Scale bars: 50 μm. (B) Quantitative analysis of 8-OHdG in lung tissue sections as described in the Methods section. (C) Representative images of TdT-mediated dUTP Nick-End Labeling (TUNEL) staining showing apoptosis in lung tissues. Scale bars: 50 μm. (D) Quantitative analysis of TUNEL in lung tissue sections as described in the Methods section. n=6, ^*^ *p* ⩽ 0.05, ^**^ *p* ⩽ 0.01, ^***^ *p* ⩽ 0.001.

### Changes in inflammatory response in lung tissue during the recovery period after exposure to HBO

Compared to 0 hours, the mRNA expression of Il6, Ccl2, and Ccl3 significantly increased at 12 hours (Il6: 4.080 ± 0.5069 vs 1.000 ± 0.1030, *p* < 0.0001; Ccl2: 4.261 ± 1.363 vs 2.521 ± 0.2703, *p* = 0.0023; Ccl3: 2.442 ± 1.267 vs 0.9443 ± 0.2209, *p* = 0.0057) (Figure 6 A, B, and C). Cxcl5 showed no significant change at 0 hours, but significantly increased after 12 hours (2.107 ± 0.6638 vs 1.000 ± 0.1221, *p* = 0.0462) (Figure 6 D). Compared to 0 hours and 12 hours, IL-6 significantly decreased after 24 hours (1.196 ± 0.3273 vs 2.287 ± 0.6055, 1.196 ± 0.3273 vs 4.080 ± 0.5069, *p* < 0.01) (Figure 3 A). Compared to 0 hours, the expression of Cxcl10 also significantly decreased after 12 and 24 hours (1.331 ± 0.2962, 1.507 ± 0.2539 vs 2.981 ± 0.7037, *p* < 0.0001) (Figure 3 E). However, the expression level of Ccl3 at 24 hours showed no difference compared to the air control group (1.573 ± 0.4477 vs 1.000 ± 0.1069, *p* = 0.4823). These results indicate that the inflammatory response continues to develop after the end of high-pressure oxygen exposure, reaching its peak at 12 hours and recovering after 24 hours.

**Fig. 6.**
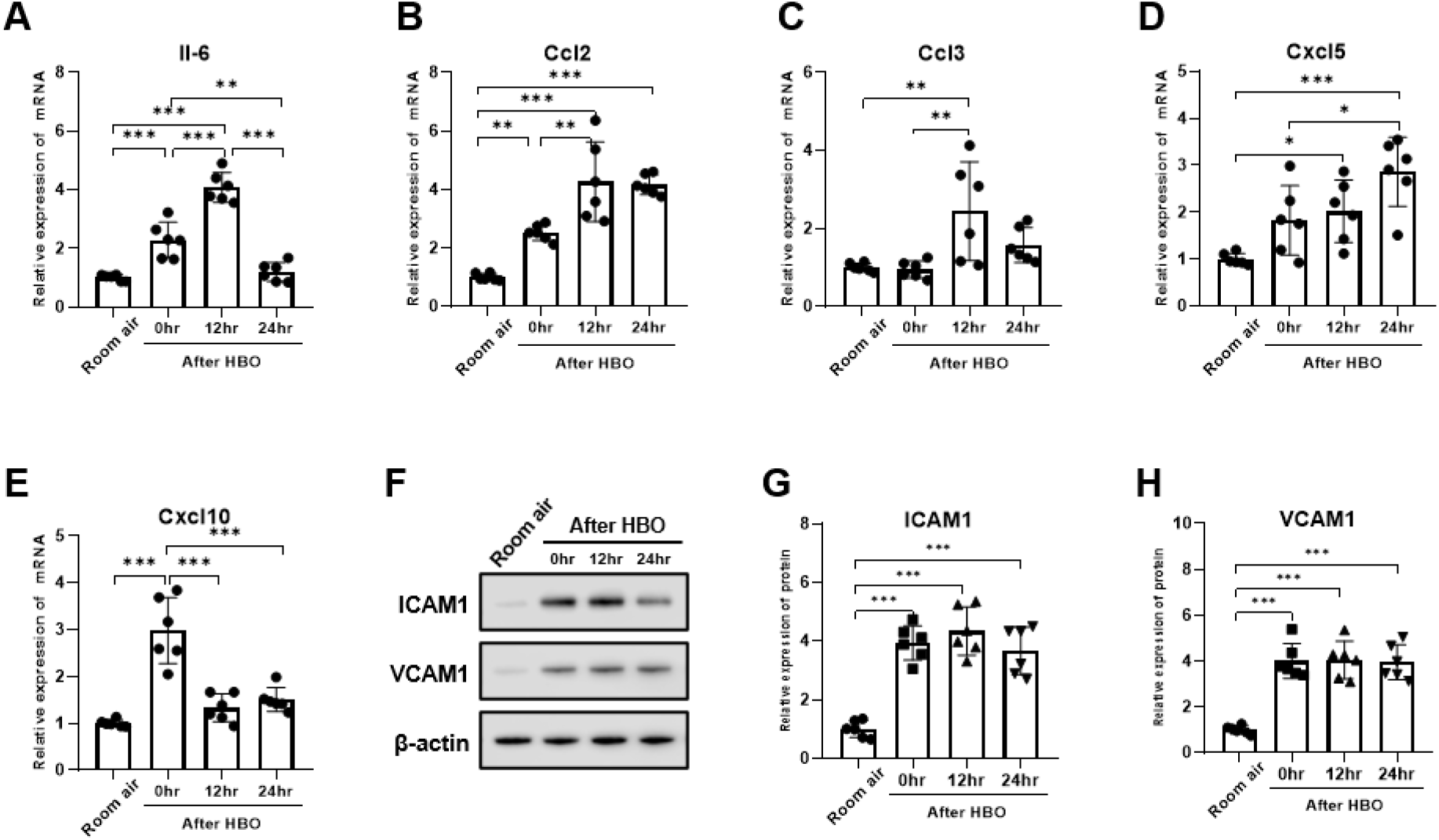
Inflammatory response in lung tissue during the recovery period after exposure to HBO. (A-E) Expression of inflammatory cytokines and chemokines in lung tissues detected by qPCR as fold change. (F) Western blot analysis of ICAM1 and VCAM1 in mouse lung samples; β-actin used as the loading control. (G, H) Quantitative graph of western blot analysis. n=6, ^*^ *p* ⩽0.05, ^**^ *p* ⩽ 0.01, ^***^ *p* ⩽ 0.001.

Western blot showed that the expression of the ICAM1 and VCAM1 proteins after HBO groups were significantly higher than the air control group (ICAM1: 3.938 ± 0.587, 4.337 ± 0.818, 3.663 ± 0.814 vs. 1.000 ± 0.289; *p* < 0.0001; VCAM1: 4.007 ±0.758, 4.052 ± 0.819, 3.953 ± 0.764 vs. 1.000 ± 0.179; *p* < 0.0001). However, during the recovery period at 0, 12, and 24 hours after exposure to HBO, there was no significant change in the expression of ICAM1 and VCAM1 proteins in lung tissue (Figure 6 F, G and H).

## DISCUSSION

This study explores the immediate impact of different HBO exposure times on lung tissue, as well as the natural progression and recovery of lung tissue after HBO exposure. The results showed that under 2ATA of HBO, there was no significant lung damage when the exposure time was less than 4 hours. However, when the exposure time exceeded 4 hours, endothelial damage in the lungs appeared before other damages, and exposure of more than 6 hours led to significant lung damage. Histological features of HBO-ALI included dilation of respiratory bronchioles and alveoli, atelectasis, inflammatory cell infiltration, and hemorrhage. Further research found that HBO exposure of more than 6 hours led to endothelial damage in lung tissue, increased vascular permeability, oxidative DNA damage in cells, and a significant up-regulation of inflammatory and chemotactic factors (such as Il6, Ccl2, Ccl3, Cxcl5, and Cxcl10). Additionally, we observed that 6 hours after exposure to 2ATA HBO, the inflammatory response and endothelial damage in lung tissue continued to progress after leaving the high oxygen environment, peaking at 12 hours and recovering after 24 hours. Twenty-four hours after leaving the high oxygen environment, the pathological changes in lung tissue, oxidative DNA damage, and the expression of inflammatory and chemotactic factors significantly decreased. We believe that in lung damage caused by HBO, the lung vascular endothelium is the most sensitive to damage and the slowest to recover. These results provide a basis for the prevention and treatment of acute lung injury complications caused by HBO.

Our research findings further substantiate that pulmonary vascular endothelial cells are the most sensitive to HBO exposure. A significant increase in EB leakage was observed just 4 hours after HBO exposure, suggesting damage to endothelial gap junctions. Western blot results indicate that ICAM1 and VCAM1 in lung tissue show an upward trend as early as 4 hours after HBO exposure, with a significant increase at 6-8 hours, preceding the expression changes of inflammatory factors. ICAM-1 and VCAM-1, as lung adhesion molecules, bind to the endothelial surface and are involved in leukocyte infiltration through the ECs and into the lungs; their elevation is considered an early event in endothelial dysfunction. These results further suggest that pulmonary vascular endothelial cells are the earliest to respond to oxygen toxicity. The damaged endothelium can increase the retention of platelets and neutrophils in the pulmonary microvasculature, promoting the subsequent occurrence and development of inflammatory responses(20, 21).

What is interesting is that we also observed that endothelial cells were the last to recover after detachment from a high-oxygen environment. Our results suggest that 24 hours after cessation of HBO, lung tissue damage scores and the expression of inflammatory factors had essentially returned to baseline, but Evans Blue (EB) leakage was still significantly increased, indicating that the intercellular junctions of the pulmonary vascular endothelium were still in a damaged state. After damage to the alveolar microvascular barrier function, the restoration of the gas exchange barrier requires multiple steps, including the proliferation of quiescent endothelial cells to replace damaged and apoptotic cells, as well as the re-establishment of adherens junctions (AJ) between microvascular endothelial cells to form a new barrier (22). Therefore, this may be the reason why pulmonary vascular endothelial permeability remains impaired after detachment from a high-oxygen environment. This research result is also consistent with clinical changes. Currently, DL_CO_ (lung diffusion capacity for carbon monoxide) is considered a sensitive index of complete recovery from pulmonary oxygen toxicity. Recovery of DLCO occurs slowly and may take several months (23). DL_CO_ is an indicator of gas diffusion function and is a sensitive indicator of damage to the blood-gas barrier. Our study suggests that the delayed recovery of lung microvascular endothelial damage might be one of the reasons for the slow recovery of DLCO following exposure to high oxygen.

The study not only examined the onset of ALI damage due to 2ATA HBO exposure but also observed the outcome of pulmonary injury after leaving the high oxygen environment. ALI induced by hyperoxia may be due to direct damage from excess ROS production, or by exacerbating inflammation (24). Our findings indicate that HBO exposure significantly increases the expression levels of inflammatory and chemotactic cytokines, including Il6, Ccl2, Ccl3, and Cxcl10, consistent with previous research on normobaric hyperoxia-induced ALI (15-18). However, under hyperbaric oxygen, significant increases in these factors are seen at 6 hours, whereas it takes at least 24 hours under normobaric conditions, with histological damage appearing at 48 hours (25). These results confirm that HBO-induced ALI requires a shorter exposure time and develops faster than normobaric oxygen. Furthermore, after leaving the HBO environment, lung tissue damage continues to progress, with inflammation persisting for at least 12 hours and beginning to recover after 24 hours. Inflammatory cytokines remained elevated at 12 hours after leaving the hyperoxic environment and began to recover at 24 hours, including Il6, Ccl2, Ccl3, Cxcl5, and Cxcl10. Cxcl5 may relate to delayed damage or repair in HBO-ALI. We observed that Cxcl5 did not change significantly immediately after 6 hours of HBO exposure but increased markedly at 12 and 24 hours post-exposure. Produced by resident lung cells (such as pulmonary epithelial cells and platelets during lung inflammation) (26), Cxcl5 plays a key role in controlling chemokine clearance (27) and can negatively regulate neutrophil influx into the lungs by affecting the concentration of other chemokines in the bloodstream (27-29). We hypothesize that Cxcl5 may suppress further inflammation in the HBO-ALI inflammatory response. Further work is needed to confirm this.

Excessive ROS from high oxygen levels directly damage cells, including DNA damage and apoptosis induction. We observed significant DNA oxidative damage in lung tissue exposed to HBO. Results indicate that DNA damage in lung tissue significantly increases after 6 hours of HBO exposure, which is markedly faster than lung damage caused by normobaric high oxygen. Nagato et al. (25) detected oxidative stress in lung tissue after exposing mice to normobaric high oxygen (100% oxygen) for 12 hours. These findings suggest again that lung damage induced by HBO progresses quicker than normobaric oxygen, consistent with our previous research (30).

The recent study did not find that HBO exposure induced significant apoptosis in lung tissue cells. In lung injury caused by normobaric hyperoxia, the apoptosis of pulmonary epithelial cells is considered to be an important characteristic of hyperoxia-induced lung injury (31, 32), but there is still controversy over cell damage caused by HBO. According to current research, whether apoptosis is induced by HBO is related to the extent of DNA damage. Schönrock et al. found that hyperbaric oxygen exposure significantly induced oxidative stress and DNA damage in osteoblast lineage cells (primary human osteoblasts, HOBs, and the osteogenic tumor cell line SAOS-2), without a significant increase in apoptosis (33). Stefan (34) and others exposed Jurkat-T-cells to environmental air or oxygen at 1-3ATA and found that HBO induced apoptosis in the cells through the mitochondrial pathway. The human promyelocytic cell line HL-60 (35), the T-lymphoblastic cell line JURKAT, and the B-lymphoblastic cell line CCRF-SB (36) were also induced to undergo apoptosis by HBO. It is currently believed that the mode of high-level oxidative stress caused by HBO exposure affects the induction of apoptosis in specific cells. HBO-induced DNA damage is mainly related to the formation of single-strand breaks (single-strand break/single-stranded DNA Damage, SSB) rather than double-strand breaks (double-strand break/double-stranded DNA damage, DSB), and such damage can be effectively repaired. The triggered repair and adaptation patterns can prevent long-term DNA damage (33), thereby avoiding the occurrence of cellular apoptosis. The exposure time for hyperoxia-induced apoptosis in lung tissue cells is often as long as 24 hours (31) or even longer (32), while our HBO intervention lasted no more than 8 hours. Therefore, we speculate that short-term HBO exposure, although it quickly results in DNA damage, is the easily repairable SSB. This damage can be effectively repaired and triggers adaptive mechanisms such as the increased expression of Hemeoxygenase-1 (HO-1) (known to increase in oxidative stress and have an anti-apoptotic effect (37)), thus preventing more severe programmed cell death (apoptosis).

### Perspectives and Significance

In summary, our study systematically observed the progression and outcomes of ALI induced by HBO. The degree of lung injury caused by HBO increased in a time-dependent manner, with pulmonary tissue damage occurring after 4 hours of exposure to 2ATA HBO. Pulmonary vascular endothelial cells were the most sensitive to high oxygen levels, sustaining the earliest damage and also recovering the slowest. The lung injury caused by 6 hours of exposure to 2ATA HBO was mainly characterized by vascular endothelial and inflammatory damage. Inflammatory responses in lung tissue peaked 12 hours after leaving the high-oxygen environment and began to recover after 24 hours, with no significant cellular apoptosis observed. The results of this study provide experimental data for understanding the development and outcomes of acute lung injury caused by HBO.

## DATA AVAILABILITY

The data that support the findings of this study are available from the corresponding author upon reasonable request.

## AUTHOR CONTRIBUTIONS

Xiaochen Bao and Guangxu Xu conceived and designed research; Shu Wang and Zhi Li performed experiments; Shu Wang and Zhi Li analyzed data; Xiaochen Bao and Shu Wang interpreted results of experiments; Shu Wang prepared figures; Xiaochen Bao and Shu Wang drafted manuscript; Xiaochen Bao and Shu Wang edited and revised manuscript; Xiaochen Bao, Shu Wang and Guangxu Xu approved final version of manuscript.

